# Immune response and resistance of clear cell renal cell carcinoma patients following immune checkpoint blockade

**DOI:** 10.1101/2023.10.05.561140

**Authors:** Guillaume Mestrallet

## Abstract

175,000 patients die because of renal cell carcinoma (RCC) each year. Clear cell renal cell carcinoma (ccRCC or KIRC) is the most frequent subtype of RCC. Current therapies include immune checkpoint inhibitors (ICB) or VEGFR tyrosine kinase inhibitors (TKIs). However, many patients did not respond to ICB and immune resistance still occurred. Immune resistance may be explained by expression of various immune checkpoints and immunosuppressive pathways in KIRC patients. Thus, it is important to identify mechanisms driving immune response and resistance following ICB. To address this question, we performed an analysis of 3 KIRC cohorts treated with 3 different ICB. Overall, 20-30% of KIRC patients respond to ICB. Responders with metastasized stage IV cancer with tumorectomy prior to anti-PD-L1 are characterized by an increase in CD4+ and CD8+ T cell infiltration, and by better antigen presentation and T cell responses (*BTN3A1, PRF1* and *CD27* genes). However, the expression of CTLA4, TIGIT and BTLA in Th1, Th17 and M2 subsets may limit complete response in responders. Importantly, non-responders patients are characterized by higher infiltration by macrophages, and by overexpression of regulatory gene (*ADORA2A*) in Th2, CD8+ T cell, M1 and M2 clusters. Targeting these pathways may help to develop combination therapies to improve KIRC patient outcomes.

## Introduction

Clear cell renal cell carcinoma (ccRCC or KIRC) is the most frequent subtype of renal cell carcinoma (RCC) (1). Around 175,000 patients die because of RCC each year (2). Current therapies include immune checkpoint inhibitors (ICB) or VEGFR tyrosine kinase inhibitors (TKIs) (3). Clinical trials were performed using various immune checkpoints, including anti-PD-1 (Nivolumab), anti-PD-L1 (Atezolizumab) and anti-CTLA4 in combination with anti-PD-1 (Nivolumab and Ipilimumab) (4–6).

Immune response at steady state or following ICB depends on antigen presentation to T cells through human leukocyte antigens (HLA) and related cell-surface molecules (MICA, MICB and ULBP) (7). T cell responses also involve the expression of genes promoting T cell proliferation, cytotoxicity and maintenance (including *BTN3A1, TNFRSF9, PRF1* and *CD27*) (8–11). On the counterpart, the expression of regulatory genes (including *ADORA2A, ARG1* and *IL-1A*) limit inflammation and maintain tissue homeostasis, but may also favor cancer development (12–14). Importantly, many patients did not respond to ICB and immune resistance still occurred (4–6). Immune resistance may be explained by expression of various immune checkpoints and immunosuppressive pathways in KIRC patients, including SIGLEC15, PD-L1, Tim-3, PDCD1, CTLA4, Lag3, PDCD1LG2, Tigit and HLA-G (1,15–17). Moreover, ICB can cause adverse events, including adrenal insufficiency and autoimmune hepatitis (18).

Thus, it is important to perform meta-analysis of KIRC patient cohorts to identify mechanisms driving immune response and resistance to different therapies. Meta-analysis of cohorts data may help to identify the best targets or combination therapies to improve patient outcomes and understand the resistance mechanisms that caused the non approval of atezolizumab in KIRC.

## Material and methods

### RNAseq datasets and selection of cohorts

Patient cohort was selected using the CRI iAtlas Portal (19). We selected the following RNAseq datasets for KIRC metastatic patients : Choueiri 2016 - KIRC, PD-1 (5), IMmotion150 - KIRC, PD-L1 (4) and Miao 2018 - KIRC, PD-1 +/-CTLA4, PD-L1 (6). We used the following group filters : Responder, Drug, Target and modified Response Evaluation Criteria in Solid Tumors (mRECIST Response). Responders are defined as patients with mRECIST of Partial Response or Complete Response, whereas Non-Responders are those with Progressive Disease or Stable Disease. Then, we used the ICI Analysis Modules. The current version of the iAtlas Portal was built in R using code hosted at https://github.com/CRI-iAtlas/iatlas-app. Assayed samples were collected prior to immunotherapy.

### Clinical description of patients

Datasets are described in **Table 1**. The Choueiri 2016 dataset contains 16 samples of 16 patients, treated with Nivolumab, an anti-PD-1. The IMmotion150 dataset contains 263 samples of 263 patients with metastasized stage IV cancer, with 89 with only tumorectomy and 174 with tumorectomy prior to Atezolizumab, an anti-PD-L1. The Miao 2018 dataset contains 17 samples of 17 patients, with 2 treated with Atezolizumab, 11 treated with Nivolumab and 4 treated with both Nivolumab and Ipilimumab, an anti-CTLA4.

**Table 1).** Clinical description of KIRC patients treated with different ICB. The Choueiri 2016 dataset contains 16 samples of 16 patients, treated with Nivolumab, an anti-PD-1. The IMmotion150 dataset contains 263 samples of 263 patients, with 89 untreated and 174 treated with Atezolizumab, an anti-PD-L1. The Miao 2018 dataset contains 17 samples of 17 patients, with 2 treated with Atezolizumab, 11 treated with Nivolumab and 4 treated with both Nivolumab and Ipilimumab, an anti-CTLA4.

| Dataset | Drug | Size | Characteristics | Target |
| --- | --- | --- | --- | --- |
| Choueiri 2016 - KIRC, PD-1 | Nivolumab | 16 | Treatment with Nivolumab | PD1 |
| IMmotion150 - KIRC, PD-L1 | Atezolizumab | 174 | Treatment with Atezolizumab | PD-L1 |
| IMmotion150 - KIRC, PD-L1 | None | 89 | No ICI therapy administered | None |
| Miao 2018 - KIRC, PD-1 +/- CTLA4, PD-L1 | Atezolizumab | 2 | Treatment with Atezolizumab | PD-L1 |
| Miao 2018 - KIRC, PD-1 +/- CTLA4, PD-L1 | Ipilimumab and Nivolumab | 4 | Combination therapy with Ipilimumab and Nivolumab | CTLA4 and PD1 |
| Miao 2018 - KIRC, PD-1 +/- CTLA4, PD-L1 | Nivolumab | 11 | Treatment with Nivolumab | PD1 |

### Immune landscape of cancer in iAtlas and immune subsets classification

The initial release of iAtlas provided a resource to complement analysis results from The Cancer Genome Atlas (TCGA) Research Network on the TCGA data set comprising over 10,000 tumor samples and 33 tumor types (“The Immune Landscape of Cancer”; here referred to as “Immune Landscape”) (20). This study identified six immune subtypes that span cancer tissue types and molecular subtypes, and found that these subtypes differ by somatic aberrations, microenvironment, and survival. Per-sample characterizations included total lymphocytic infiltrate (from DNA methylation as well as H&E imaging data), estimated cell type fractions, immune gene signature expression, MHC/HLA type and expression, antigen presentation machinery, T cell and B cell receptor repertoire inference, viral/microbial characterization, associations with pathway disruption and activity, and other analysis results. The Immune Landscape manuscript reported on the most novel and potentially therapeutically salient statistical associations between these immune subtypes and the results of the immune characterization (20). C1 (Wound Healing) had elevated expression of angiogenic genes, a high proliferation rate, and a Th2 cell bias to the adaptive immune infiltrate. C2 (IFN-γ Dominant) had the highest M1/M2 macrophage polarization, a strong CD8 signal and, together with C6, the greatest TCR diversity. C2 also showed a high proliferation rate, which may override an evolving Type I immune response. C3 (Inflammatory) was defined by elevated Th17 and Th1 genes, low to moderate tumor cell proliferation, and, along with C5, lower levels of aneuploidy and overall somatic copy number alterations than the other subtypes. C4 (Lymphocyte Depleted) displayed a more prominent macrophage signature, with Th1 suppressed and a high M2 response. C5 (Immunologically Quiet), exhibited the lowest lymphocyte, and highest macrophage responses, dominated by M2 macrophages. IDH mutations were enriched in C5 over C4 (80% of IDH mutations), suggesting an association of IDH-mutations with favorable immune composition. Indeed, IDH-mutations are associated with TME composition and decrease leukocyte chemotaxis, leading to fewer tumor-associated immune cells and better outcome. Finally, C6 (TGF-β Dominant) displayed the highest TGF-β signature and a high lymphocytic infiltrate with an even distribution of Type I and Type II T cells.

### Statistics

Statistical significance of the observed differences was determined using the independent t-Test and Wilcoxon t-Test. All data are presented as mean±SEM. The difference was considered as significant when the p value was below 0.05. * : p<0.05 for both tests.

## Results

### Clinical responses of patients treated with different ICB

We first performed a meta-analysis of clinical responses of KIRC patients following various ICB therapies (**Table 2 and Figure 1**). The percentage of responders after ICB are the same in IMmotion150 (Atezolizumab) and Miao 2018 (2 treated with Atezolizumab, 11 treated with Nivolumab and 4 treated with both Nivolumab and Ipilimumab) datasets (29%) but lower for Choueiri 2016 (Nivolumab) study (19%). Thus, treatment with Atezolizumab or Nivolumab +/-Ipilimumab gave similar mRECIST responses. However, few patients were treated with Nivolumab + Ipilimumab, limiting a robust statistical analysis. In more detail, in the IMmotion150 cohort, we observed 9.2% with complete response, 18.4% with partial response, 37.4% with stable disease and 29.9% with progressive disease following ICB. Overall, 20-30% of KIRC patients respond to ICB.

**Table 2).** Clinical response of KIRC patients treated with different ICB. Patients are stratified using modified Response Evaluation Criteria in Solid Tumors (mRECIST Response).

| Dataset | Percentage Non-Responders (%) | Percentage Responders (%) | Size Non-Response | Size Response | Size Complete Response | Size Partial Response | Size Stable Disease | Size Progressive Disease | Size Not Available |
| --- | --- | --- | --- | --- | --- | --- | --- | --- | --- |
| Choueiri 2016 - KIRC, PD-1 | 81 | 19 | 13 | 3 | 0 | 3 | 5 | 8 | 0 |
| IMmotion 150 - KIRC, PD-L1 | 71 | 29 | 117 | 48 | 16 | 32 | 65 | 52 | 9 |
| Miao 2018 - KIRC, PD-1 +/- CTLA4, PD-L1 | 71 | 29 | 12 | 5 | 1 | 4 | 4 | 8 | 0 |

**Figure 1).**
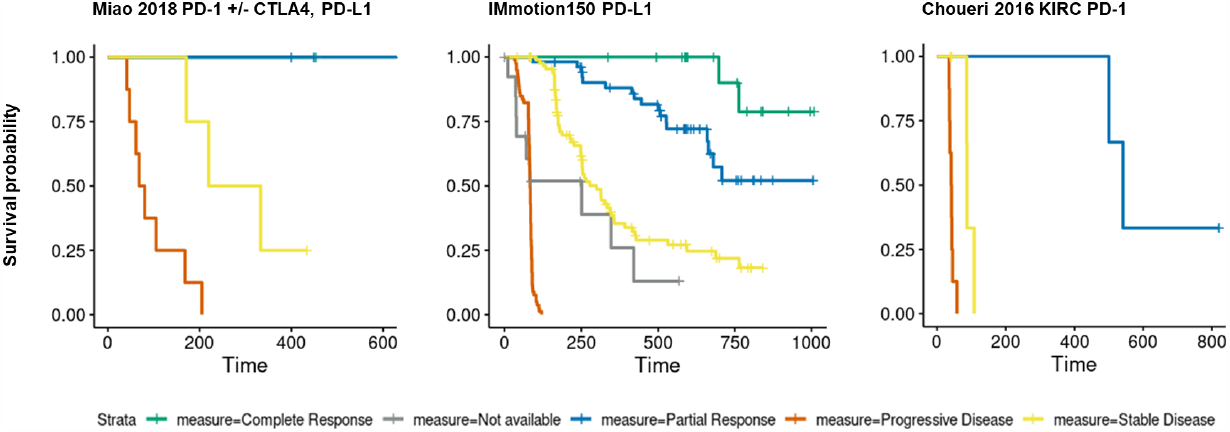
Response following anti-PD-L1 / anti-PD-1 / anti-CTLA4 + antiPD-1 therapy in KIRC patients. The Choueiri 2016 dataset contains 16 samples of 16 patients, treated with Nivolumab, an anti-PD-1. The IMmotion150 dataset contains 263 samples of 263 patients, with 89 untreated and 174 treated with Atezolizumab, an anti-PD-L1. The Miao 2018 dataset contains 17 samples of 17 patients, with 2 treated with Atezolizumab, 11 treated with Nivolumab and 4 treated with both Nivolumab and Ipilimumab, an anti-CTLA4. Patients are stratified using modified Response Evaluation Criteria in Solid Tumors (mRECIST Response).

### Response and immune infiltration following anti-PD-L1 therapy in KIRC patients

We investigated if response to anti-PD-L1 therapy correlated with changes in immune infiltration. We performed RNAseq analysis in the IMmotion150 cohort to have robust statistical analysis (**Figure 2**). We confirmed an increase in survival probability after Atezolizumab in responders with metastasized stage IV cancer with tumorectomy prior to anti-PD-L1 (**Figure 2 A**). We observed more lymphocytes and in more details more CD8+ and CD4+ T cell infiltration in responders (**Figure 2 B**). However we observed less NK cells resting, DC cell resting and macrophage infiltration in responders. We have not observed differences in other immune subsets infiltration (including B cells, Eosinophils, Mast cells, Neutrophils, Monocytes, Plasma cells and Tregs). Overall, ICB increased T-cell infiltration in responders. However, immune resistance to ICB in patients with metastasized stage IV cancer with tumorectomy prior to anti-PD-L1 may be triggered by an increase in macrophage infiltration.

**Figure 2).**
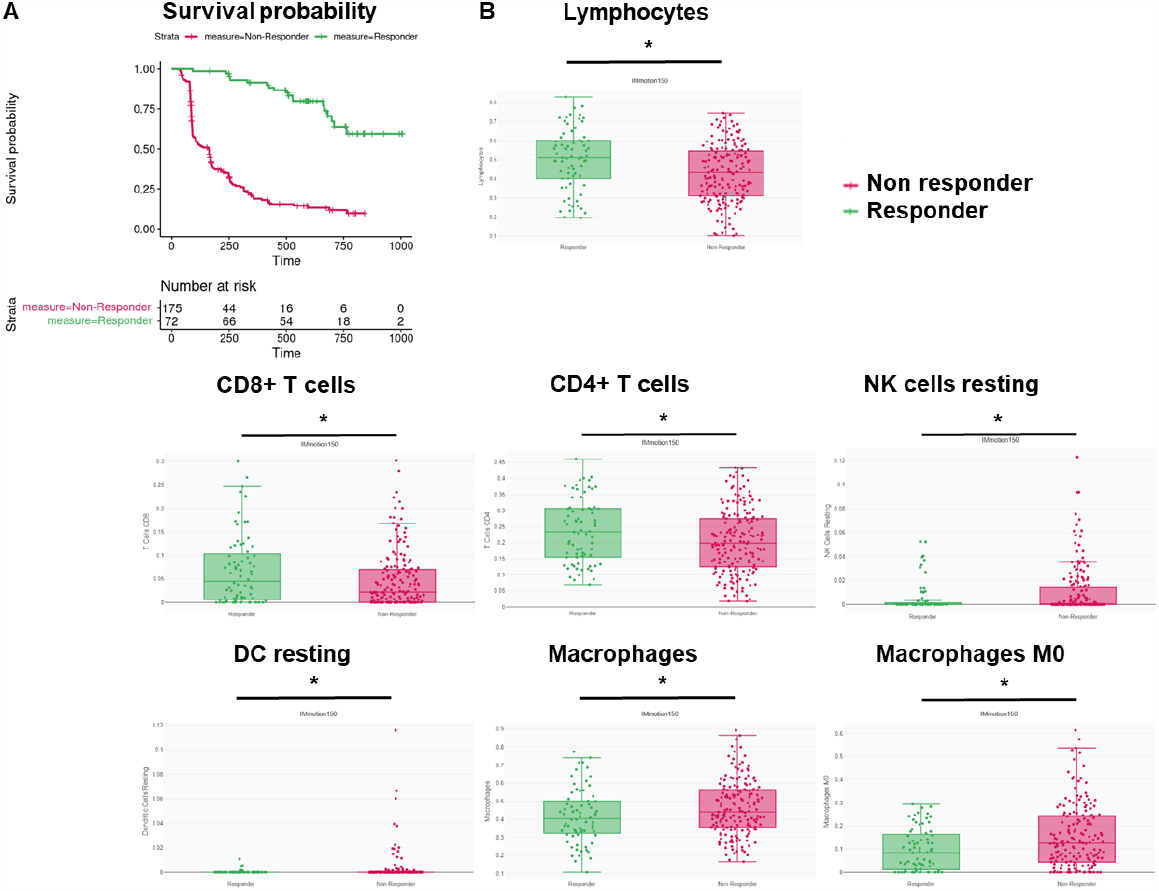
Response and immune infiltration following anti-PD-L1 therapy in KIRC patients. A Survival probability in IMmotion150 cohort after Atezolizumab therapy in responders and non-responders. **B** Immune infiltration measured by RNAseq in IMmotion150 cohort after Atezolizumab therapy in responders and non-responders. Responder = Patient with mRECIST of Partial Response or Complete Response. N = 247. Independent t-Test and Wilcoxon t-Test.

### Immune gene expression following anti-PD-L1 therapy in KIRC patients

We investigated if response to anti-PD-L1 therapy correlated with changes in immune gene expression in tumors. We performed RNAseq analysis in the IMmotion150 cohort to have robust statistical analysis (**Figure 3**). We observed more antigen presentation genes (*HLA-DPB1, HLA-DPA1* and *MICB*) expression in responders (**Figure 3 A**). We also observed more T cell proliferation (*BTN3A1*), cytotoxic (*PRF1*) and maintenance (*CD27*) genes in responders (**Figure 3 B**). However, we observed an increase in some immune checkpoint genes (*TIGIT* and *CTLA4*) and *BLTA* immunosuppressive gene in responders (**Figure 3 C/D**). Importantly, we observed less immunosuppressive (*ADORA2A*) gene expression in responders (**Figure 3 D**). We have not observed differences in expression of other immunoregulatory genes and immune checkpoints (including TIM3, LAG3, PD-L1, PD-L2, EDNRB, TLR4, IDO1, ARG1, VSIR, BTN3A, CCL5, CD40, TNFRSF, CD28, CD80, ICOS, VTCN1, TNFSF, CD70, CX3CL1, CXCL10, CXCL9, ENTPD1, GXMA, HMGB1, ICAM1, ICOSLG, VEGF, KIR, IFN/TNF/TGFB genes, interleukines, MICA and other HLA genes). Overall, ICB responders with metastasized stage IV cancer with tumorectomy prior to anti-PD-L1 were characterized by better antigen presentation and T cell responses.

**Figure 3).**
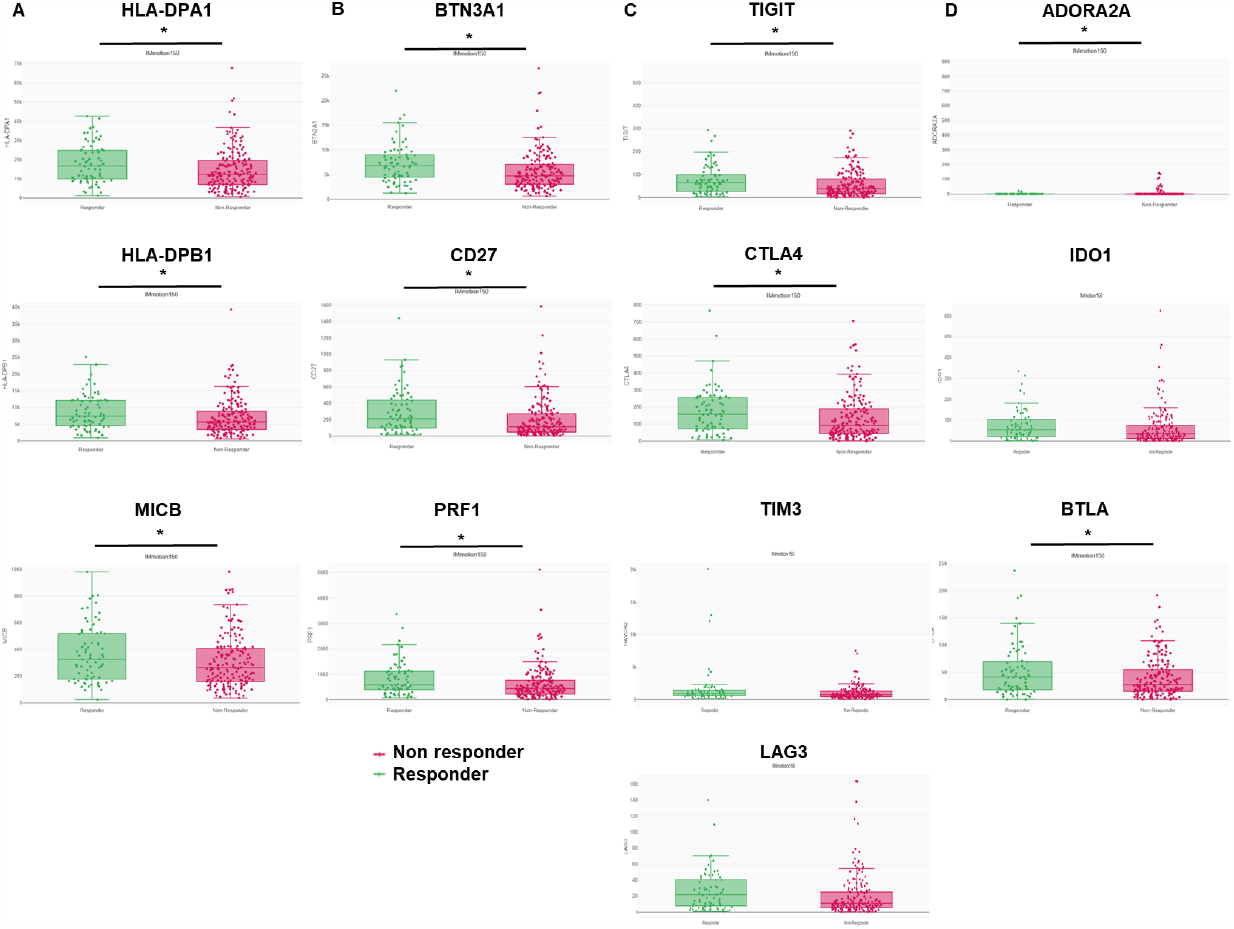
Immune gene expression following anti-PD-L1 therapy in KIRC patients. A Antigen presentation genes expression in IMmotion150 cohort after Atezolizumab therapy in responders and non-responders. **B** T cell proliferation, cytotoxic and maintenance genes expression in IMmotion150 cohort after Atezolizumab therapy in responders and non-responders. **C** Immune checkpoints gene expression in IMmotion150 cohort after Atezolizumab therapy in responders and non-responders. **D** Immunosuppressive and regulatory genes expression in IMmotion150 cohort after Atezolizumab therapy in responders and non-responders. Responder = Patient with mRECIST of Partial Response or Complete Response. N = 247. Independent t-Test and Wilcoxon t-Test.

### Immune subsets involved in resistance to anti-PD-L1 therapy in KIRC patients

We observed that TIGIT and CTLA4 checkpoints and BTLA were expressed in C3, C6 and C4 immune clusters in both responders and non-responders (**Figure 4**). We also observed that ADORA2A was expressed in the C3 cluster in responders and non-responders (**Figure 4**). C3 (Inflammatory) was defined by elevated Th17 and Th1 genes (20). C4 (Lymphocyte Depleted) displayed a more prominent macrophage signature, with Th1 suppressed and a high M2 response. C6 (TGF-β Dominant) displayed the highest TGF-β signature and a high lymphocytic infiltrate with an even distribution of Type I and Type II T cells. Interestingly, TIGIT and CTLA4 checkpoints and BTLA were also expressed in C1 and C2 immune clusters in non-responders, and ADORA2A was also expressed in C2, C4 and C6 clusters in non-responders (**Figure 4**). C1 (Wound Healing) had elevated expression of angiogenic genes, a high proliferation rate, and a Th2 cell bias to the adaptive immune infiltrate. C2 (IFN-γ Dominant) had the highest M1/M2 macrophage polarization, a strong CD8 signal and, together with C6, the greatest TCR diversity (20). Overall, immune resistance in responders may be explained by *TIGIT* and *CTLA4* checkpoints and *BTLA* overexpression in Th1, Th17 and M2 subsets. Immune resistance in non-responders involves ADORA2A expression in CD8+ T cell, M1 and M2 clusters compared to responders.

**Figure 4).**
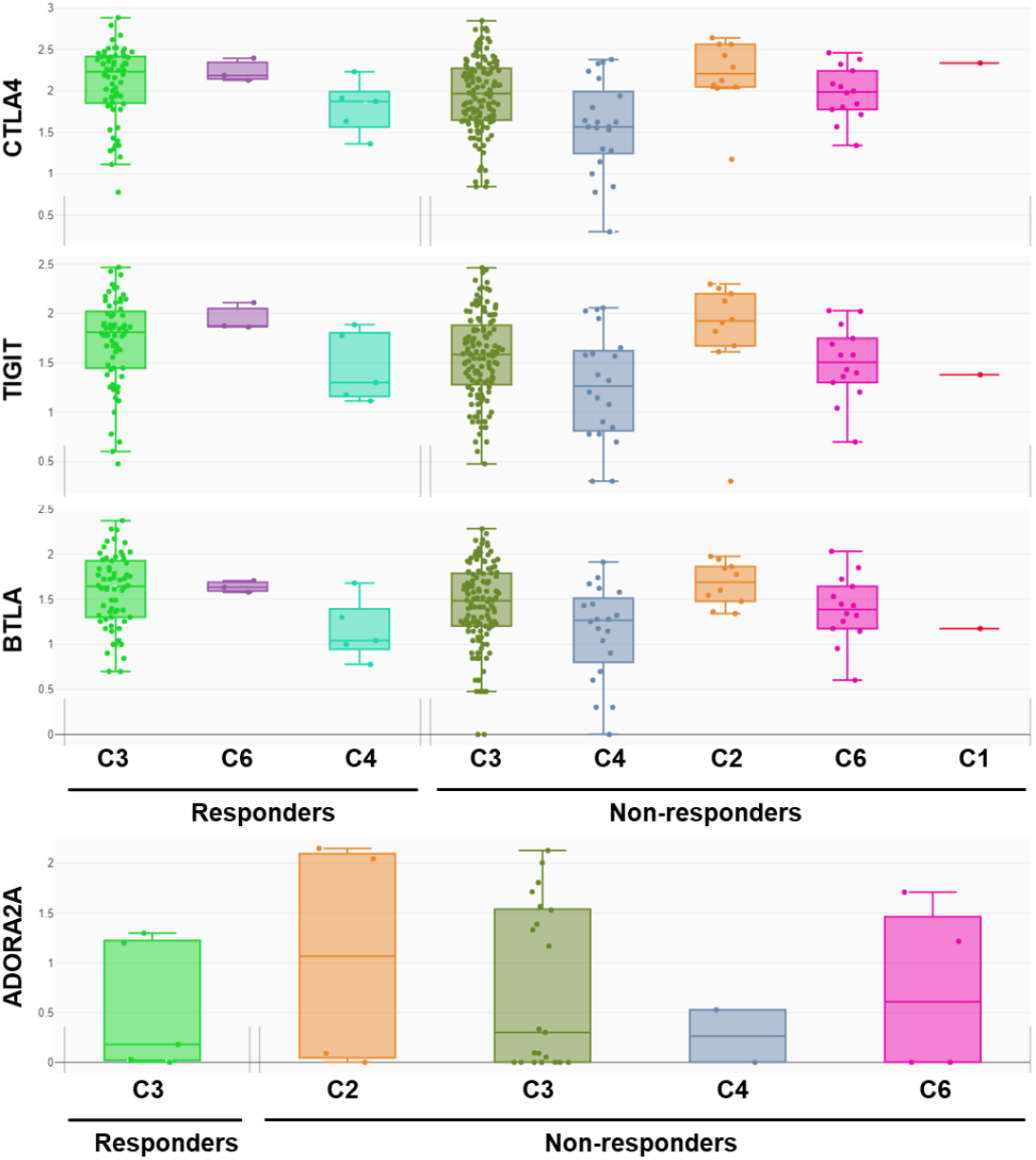
Immunosuppressive gene expression in 6 immune subtype clusters in responders and non-responders to ICB. The iAtlas provided a resource to complement analysis results from TCGA Research Network on the TCGA data set comprising over 10,000 tumor samples and 33 tumor types (“The Immune Landscape of Cancer”) (20). This study identified six immune subtypes that span cancer tissue types and molecular subtypes, and found that these subtypes differ by somatic aberrations, microenvironment, and survival. C1 (Wound Healing) had elevated expression of angiogenic genes, a high proliferation rate, and a Th2 cell bias to the adaptive immune infiltrate. C2 (IFN-γ Dominant) had the highest M1/M2 macrophage polarization, a strong CD8 signal and, together with C6, the greatest TCR diversity. C2 also showed a high proliferation rate, which may override an evolving Type I immune response. C3 (Inflammatory) was defined by elevated Th17 and Th1 genes, low to moderate tumor cell proliferation, and, along with C5, lower levels of aneuploidy and overall somatic copy number alterations than the other subtypes. C4 (Lymphocyte Depleted) displayed a more prominent macrophage signature, with Th1 suppressed and a high M2 response. C5 (Immunologically Quiet), exhibited the lowest lymphocyte, and highest macrophage responses, dominated by M2 macrophages. IDH mutations were enriched in C5 over C4 (80% of IDH mutations), suggesting an association of IDH-mutations with favorable immune composition. Indeed, IDH-mutations are associated with TME composition and decrease leukocyte chemotaxis, leading to fewer tumor-associated immune cells and better outcome. Finally, C6 (TGF-β Dominant) displayed the highest TGF-β signature and a high lymphocytic infiltrate with an even distribution of Type I and Type II T cells.

## Discussion

By performing meta-analysis on 3 KIRC cohorts, we showed that 20-30% of KIRC patients respond to different ICB, including anti-PD-1 (Nivolumab), anti-PD-L1 (Atezolizumab) and anti-CTLA4 in combination with anti-PD-1 (Nivolumab and Ipilimumab). Responders with metastasized stage IV cancer with tumorectomy prior to anti-PD-L1 blockade are characterized by an increase in lymphocytes and more specifically T cell infiltration following ICB. They are also characterized by better antigen presentation (HLA-DPA1/DPB1, MICB) and T cell responses (BTN3A1, CD27, PRF1). Of note, the CD27 axis was also identified in a transcriptome KIRC study (21).

However, some immunosuppressive pathways, including TIGIT and CTLA4 checkpoints and BTLA are overexpressed in responders. It may limit the response of these patients following ICB. These results are in line with previous studies, showing that high expression of CTLA4 was associated with poor overall survival, progression-free survival, and disease-free survival of KIRC patients (1). Here, we show that the expression of these resistance pathways in responders involve the Th1, Th17 and M2 subsets. Combination therapy with Ipilimumab and Nivolumab to target PD-1 and CTLA4 is promising, but more replicates are needed in clinical settings to assess a potential interest. Indeed, only 4 patients were treated with Nivolumab + Ipilimumab, limiting a robust statistical analysis (6).

Importantly, 70-80% of KIRC patients with metastasized stage IV cancer with tumorectomy prior to anti-PD-L1 did not respond to ICB. Non-responders patients are characterized by higher infiltration by myeloid subsets, including DC cell resting and macrophages. This is in line with studies showing that immune resistance following ICB may be mediated by myeloid subsets (22–24). Here, we show that non-responders to ICB are characterized by an increase in immunoregulatory ADORA2A gene expression in CD8+ T cell, M1 and M2 clusters. Targeting ADORA2A was investigated in mouse tumor models and CART cell assays and limited tumor growth (25,26).

Overall, it may be useful to test drugs targeting multiple immune checkpoint (*TIGIT* and *CTLA4*) and *BLTA* in partial KIRC responders to limit resistance to ICB monotherapy. In addition, we may test drugs targeting macrophages and immunoregulatory pathways (*ADORA2A*) in non-responders to make them sensitive to ICB. This may be done first in vitro, using patient tumor-derived spheroids co-cultured with autologous immune cells, as done with anti-PD-1, then in vivo in mice models and next if validated in KIRC patients (3). Thus, meta-analysis of cohort data with more patients treated with anti-PD-1 + anti-CTLA4 or anti-Tigit may help to assess the best targets or combination therapies to improve KIRC patient outcomes.

## References

1. Liao G, Wang P, Wang Y. Identification of the Prognosis Value and Potential Mechanism of Immune Checkpoints in Renal Clear Cell Carcinoma Microenvironment. Front Oncol (2021) 11: https://www.frontiersin.org/articles/10.3389/fonc.2021.720125 x[Accessed October 12, 2022]

2. Bray F, Ferlay J, Soerjomataram I, Siegel RL, Torre LA, Jemal A. Global cancer statistics 2018: GLOBOCAN estimates of incidence and mortality worldwide for 36 cancers in 185 countries. CA Cancer J Clin (2018) 68:394–424. doi: 10.3322/caac.21492

3. Lugand L, Mestrallet G, Laboureur R, Dumont C, Bouhidel F, Djouadou M, Masson-Lecomte A, Desgrandchamps F, Culine S, Carosella ED, et al. Methods for Establishing a Renal Cell Carcinoma Tumor Spheroid Model With Immune Infiltration for Immunotherapeutic Studies. Front Oncol (2022) 12: https://www.frontiersin.org/articles/10.3389/fonc.2022.898732 x[Accessed July 28, 2022]

4. McDermott DF, Huseni MA, Atkins MB, Motzer RJ, Rini BI, Escudier B, Fong L, Joseph RW, Pal SK, Reeves JA, et al. Clinical activity and molecular correlates of response to atezolizumab alone or in combination with bevacizumab versus sunitinib in renal cell carcinoma. Nat Med (2018) 24:749–757. doi: 10.1038/s41591-018-0053-3

5. Choueiri TK, Powles T, Burotto M, Escudier B, Bourlon MT, Zurawski B, Oyervides Juárez VM, Hsieh JJ, Basso U, Shah AY, et al. Nivolumab plus Cabozantinib versus Sunitinib for Advanced Renal-Cell Carcinoma. N Engl J Med (2021) 384:829–841. doi: 10.1056/NEJMoa2026982

6. Miao X, Xu R, Fan B, Chen J, Li X, Mao W, Hua S, Li B. PD-L1 reverses depigmentation in Pmel-1 vitiligo mice by increasing the abundance of Tregs in the skin. Sci Rep (2018) 8:1–6. doi: 10.1038/s41598-018-19407-w

7. Mestrallet G, Rouas-Freiss N, LeMaoult J, Fortunel NO, Martin MT. Skin Immunity and Tolerance: Focus on Epidermal Keratinocytes Expressing HLA-G. Front Immunol (2021) 12: https://www.frontiersin.org/article/10.3389/fimmu.2021.772516 [Accessed March 8, 2022]

8. Payne KK, Mine JA, Biswas S, Chaurio RA, Perales-Puchalt A, Anadon CM, Costich TL, Harro CM, Walrath J, Ming Q, et al. BTN3A1 governs antitumor responses by coordinating αβ and γδ T cells. Science (2020) 369:942–949. doi: 10.1126/science.aay2767

9. Fröhlich A, Loick S, Bawden EG, Fietz S, Dietrich J, Diekmann E, Saavedra G, Fröhlich H, Niebel D, Sirokay J, et al. Comprehensive analysis of tumor necrosis factor receptor TNFRSF9 (4-1BB) DNA methylation with regard to molecular and clinicopathological features, immune infiltrates, and response prediction to immunotherapy in melanoma. eBioMedicine (2020) 52: doi: 10.1016/j.ebiom.2020.102647

10. Fan C, Hu H, Shen Y, Wang Q, Mao Y, Ye B, Xiang M. PRF1 is a prognostic marker and correlated with immune infiltration in head and neck squamous cell carcinoma. Transl Oncol(2021) 14:101042. doi: 10.1016/j.tranon.2021.101042

11. Hendriks J, Gravestein LA, Tesselaar K, van Lier RAW, Schumacher TNM, Borst J. CD27 is required for generation and long-term maintenance of T cell immunity. Nat Immunol (2000) 1:433–440. doi: 10.1038/80877

12. Cekic C, Linden J. Adenosine A2A receptors intrinsically regulate CD8+ T cells in the tumor microenvironment. Cancer Res (2014) 74:7239–7249. doi: 10.1158/0008-5472.CAN-13-3581

13. Bronte V, Serafini P, De Santo C, Marigo I, Tosello V, Mazzoni A, Segal DM, Staib C, Lowel M, Sutter G, et al. IL-4-induced arginase 1 suppresses alloreactive T cells in tumor-bearing mice. J Immunol Baltim Md 1950 (2003) 170:270–278. doi: 10.4049/jimmunol.170.1.270

14. Mantovani A, Barajon I, Garlanda C. IL-1 and IL-1 Regulatory Pathways in Cancer Progression and Therapy. Immunol Rev (2018) 281:57–61. doi: 10.1111/imr.12614

15. Wei SC, Duffy CR, Allison JP. Fundamental Mechanisms of Immune Checkpoint Blockade Therapy. Cancer Discov (2018) 8:1069–1086. doi: 10.1158/2159-8290.CD-18-0367

16. Pardoll DM. The blockade of immune checkpoints in cancer immunotherapy. Nat Rev Cancer (2012) 12:252–264. doi: 10.1038/nrc3239

17. Rouas-Freiss N, LeMaoult J, Verine J, Tronik-Le Roux D, Culine S, Hennequin C, Desgrandchamps F, Carosella ED. Intratumor heterogeneity of immune checkpoints in primary renal cell cancer: Focus on HLA-G/ILT2/ILT4. Oncoimmunology (2017) 6:e1342023. doi: 10.1080/2162402X.2017.1342023

18. Parikh M, Bajwa P. Immune Checkpoint Inhibitors in the Treatment of Renal Cell Carcinoma. Semin Nephrol (2020) 40:76–85. doi: 10.1016/j.semnephrol.2019.12.009

19. Eddy JA, Thorsson V, Lamb AE, Gibbs DL, Heimann C, Yu JX, Chung V, Chae Y, Dang K, Vincent BG, et al. CRI iAtlas: an interactive portal for immuno-oncology research. (2020) doi: 10.12688/f1000research.25141.1

20. Thorsson V, Gibbs DL, Brown SD, Wolf D, Bortone DS, Yang T-HO, Porta-Pardo E, Gao G, Plaisier CL, Eddy JA, et al. The Immune Landscape of Cancer. Immunity (2018) 48:812–830.e14. doi: 10.1016/j.immuni.2018.03.023

21. Alchahin AM, Mei S, Tsea I, Hirz T, Kfoury Y, Dahl D, Wu C-L, Subtelny AO, Wu S, Scadden DT, et al. A transcriptional metastatic signature predicts survival in clear cell renal cell carcinoma. Nat Commun (2022) 13:5747. doi: 10.1038/s41467-022-33375-w

22. Mestrallet G, Sone K, Bhardwaj N. Strategies to overcome DC dysregulation in the tumor microenvironment. Front Immunol (2022) 13: https://www.frontiersin.org/articles/10.3389/fimmu.2022.980709 [Accessed October 12, 2022]

23. Wang L, Sfakianos JP, Beaumont KG, Akturk G, Horowitz A, Sebra RP, Farkas AM, Gnjatic S, Hake A, Izadmehr S, et al. Myeloid Cell-associated Resistance to PD-1/PD-L1 Blockade in Urothelial Cancer Revealed Through Bulk and Single-cell RNA Sequencing. Clin CancerRes Off J Am Assoc Cancer Res (2021) 27:4287–4300. doi: 10.1158/1078-0432.CCR-20-4574

24. Nebot-Bral L, Hollebecque A, Yurchenko AA, de Forceville L, Danjou M, Jouniaux J-M, Rosa RCA, Pouvelle C, Aoufouchi S, Vuagnat P, et al. Overcoming resistance to αPD-1 of MMR-deficient tumors with high tumor-induced neutrophils levels by combination of αCTLA-4 and αPD-1 blockers. J Immunother Cancer (2022) 10:e005059. doi: 10.1136/jitc-2022-005059

25. Beavis PA, Henderson MA, Giuffrida L, Mills JK, Sek K, Cross RS, Davenport AJ, John LB, Mardiana S, Slaney CY, et al. Targeting the adenosine 2A receptor enhances chimeric antigen receptor T cell efficacy. J Clin Invest (2017) 127:929–941. doi: 10.1172/JCI89455

26. Sitkovsky MV, Kjaergaard J, Lukashev D, Ohta A. Hypoxia-Adenosinergic Immunosuppression: Tumor Protection by T Regulatory Cells and Cancerous Tissue Hypoxia. Clin Cancer Res (2008) 14:5947–5952. doi: 10.1158/1078-0432.CCR-08-0229

